# Visual looming is a primitive for human emotion

**DOI:** 10.1101/2023.08.29.555380

**Authors:** Monica K. Thieu, Vladislav Ayzenberg, Stella F. Lourenco, Philip A. Kragel

## Abstract

Looming objects afford threat of collision across the animal kingdom. Defensive responses to looming and neural computations for looming detection are strikingly conserved across species. In mammals, information about rapidly approaching threats is conveyed from the retina to the midbrain superior colliculus, where variables that indicate the position and velocity of approach are computed to enable defensive behavior. Although neuroscientific theories posit that midbrain representations contribute to emotion through connectivity with distributed brain systems, it remains unknown whether a computational system for looming detection can predict both defensive behavior and phenomenal experience in humans. Here, we show that a shallow convolutional neural network based on the *Drosophila* visual system predicts defensive blinking to looming objects in infants and superior colliculus responses to optical expansion in adults. Further, the responses of the convolutional network to a broad array of naturalistic video clips predict self-reported emotion largely on the basis of subjective arousal. Our findings illustrate how motor and experiential components of human emotion relate to species-general systems for survival in unpredictable environments.

## Main Text

Emotions guide people to make sense of and react adaptively to the world around them. A hallmark of human emotion is the complexity of emotionally evocative situations and the varied ways in which they are appraised. Nevertheless, certain events consistently drive similar experiences across individuals. A spectator at a baseball game is likely to flinch in the face of an oncoming foul ball. A pedestrian might report feeling frightened after a speeding car cuts too close to them in the crosswalk. Even if emotional experience is ultimately highly personalized by a variety of developmental and cultural factors^1^, some aspects of this experience are likely built upon mechanisms that are shared across people and across phylogeny. These building blocks of emotion are considered “primitives” in the sense that they are psychologically irreducible^2^ and they have properties that are present across species^3^. Thus, to understand the nature and origins of human emotion, we must identify which features are shared across species and the means by which specific sensory inputs drive specific emotional states.

Humans are tuned to detect and react to certain classes of ancestrally relevant stimuli, and threats to survival in particular ^4,5^. Predators make up one such class of threats. For example, human observers— including infants and children—detect images of snakes faster than other objects^6–10^. Macaques also rapidly detect and learn to avoid snakes^11,12^, two behaviors thought to be implemented in subcortical pathways through the superior colliculus and pulvinar^13^. Emotional expressions make up another such class of sensory signals indicative of threat. For example, fearful facial expressions are detected more rapidly than other expressions^14^. This heightened sensitivity may be subserved by the detection of specific visual features, like widened eyes, in the amygdala via similar inputs from the pulvinar nucleus^15^. Although these findings may be taken to suggest that threats are detected through similar neural mechanisms, not all animals are as sensitive to predatory snakes, or the wide-eyed facial expressions of conspecifics, suggesting these behaviors are unlikely to be supported by neural mechanisms that are shared across species.

One type of stimulus that is generally perceived as threatening and evokes defensive behavior across species is visual looming. As an object approaches the viewer, or *looms,* it tends to block light, and its edges expand optically. Additionally, if the object is on a collision course, its edges will expand radially in the observer’s frame of reference. Rapidly approaching objects in the environment are almost invariably dangerous, like predators, or projectiles that may cause physical damage upon contact, and very few other types of environmental motion will create such a combination of visual features. Dark-shape radial expansion thus affords threat of collision to any animal that can detect it^16^.

Many species of animals show defensive responses to looming stimuli that are subserved by functionally similar neural pathways. Rapidly looming shadows elicit escape behaviors in animals including but not limited to insects, birds, rodents, and nonhuman primates^17–20^. Humans, as well, show defensive responses—when faced with physically looming objects, human infants and adults blink and flinch respectively^21–24^. Across mammals, detecting and responding to looming motion involves the superior colliculus, a midbrain structure whose neural organization and role in sensorimotor orienting is highly conserved across species^25^. Indeed, the human superior colliculus responds more strongly to looming than receding visual stimuli^26^. Information about looming is used to coordinate defensive behavior via projections to subcortical structures including the periaqueductal grey, ventral tegmental area, and the thalamus ^27^. The computations involved in detecting and responding to visually looming threats are comparable across vertebrates^28^, suggesting they may produce a “central emotion state” that is a building block of emotion ^3^. This conservation suggests models of looming detection from nonhuman animal studies can be applied to predict human responses to similar stimuli.

We hypothesized that visual looming contributes to human emotional experience via computations that are conserved across species and only require information available in the optical array ^16^. If this is the case, then a species-general neural network optimized for collision detection should predict brain activity, defensive responses to looming objects, and subjective experience in humans. Here we tested this hypothesis in three ways. First, we assessed whether representations of looming from the convolutional neural network are encoded in patterns of superior colliculus responses to dynamic videos^29^ in human adults. Second, we tested whether the convolutional neural network predicts defensive blinking to looming objects in human infants. Third, we evaluated whether representations of looming relate to valence and arousal or specific emotion categories, using the neural network to predict self-reported following exposure to naturalistic videos^30^. Through these analyses, we test which aspects of human affective experience could be modeled by a simple computational system based on algorithms implemented in the nervous system of multiple species.

### Visual looming is encoded in the human superior colliculus

To model looming, we adapted a pre-trained shallow convolutional neural network with connections constrained by the connectivity of *Drosophila* LPLC2 neurons ^33^, directly inputting the pre-trained filter for a single LPLC2 “neuron” as the kernel (Figure 1A). Unlike parametric models that use variables such as the relative rate of expansion τ (tau) and the optical variable η (eta) to compute the approach of looming objects^31,32^, the network takes sequences of optical flow as input, providing a model that can process naturalistic videos. Four channels of inputs are analyzed per frame–one for each of the cardinal directions of optical flow—and each channel is convolved with a characteristic radial outward motion filter, producing a two-dimensional spatial representation of looming. This representation is summed across units to produce a framewise estimate of collision probability over the sequence of visual inputs.

**Figure 1.**
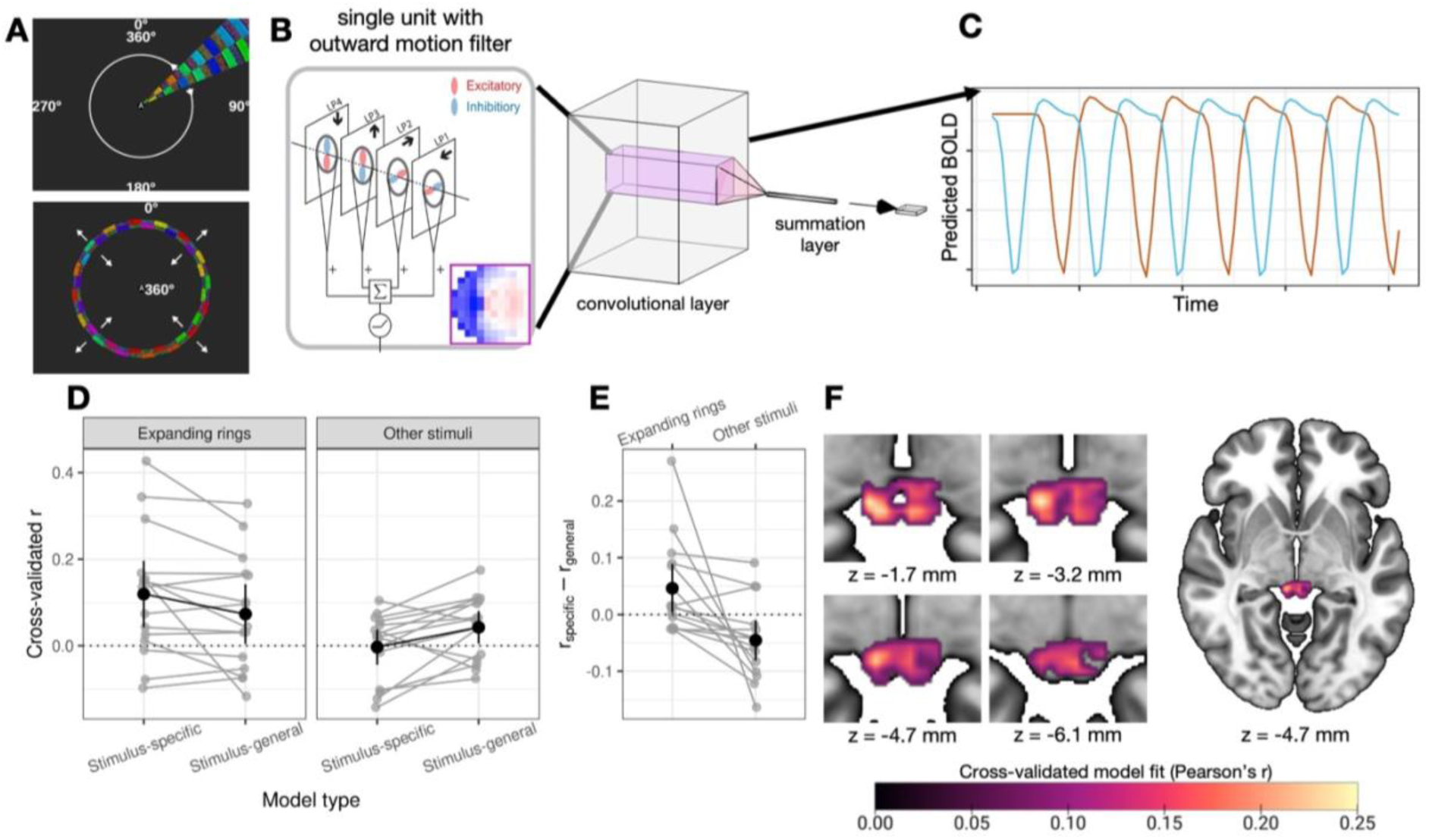
Visual looming is encoded in the human superior colliculus. **(A)** Dynamic retinotopic mapping stimuli featuring clockwise and counterclockwise sweeping wedges (top) and contracting and expanding rings (bottom) used in the fMRI experiment. **(B)** The shallow convolutional neural network originally trained to detect imminent collision. The pre-trained convolutional units (left) filter each frame for outward motion in the four cardinal directions and output a matrix of activations corresponding to the timecourse of looming motion at various points in the visual field. Panel adapted with permission from reference^33^. **(C)** Exemplar timecourses used to fit encoding models. Predictor variables are shown for a 5-cycle run of the expanding ring stimulus from two units at the center (blue) and periphery (orange) of the visual field. Units at the center tend to peak in activation early in the cycle, when the ring is in the center of the visual field, and units at the periphery tend to peak later in the cycle, when the ring has expanded. **(D)** Model performance estimated using leave-one-subject-out cross-validated Pearson’s *r* between encoding model-predicted and observed BOLD. Gray points and lines show model fit estimates for each held-out subject. Black summary points and error bars show mean ±2 standard errors across cross-validation folds. The expansion-specific model of superior colliculus activity outperforms a stimulus-general model on the same data (left subplot). **(E)** Difference in model fit between the stimulus-specific and stimulus-general encoding models for expanding rings. **(F)** Voxelwise activity explained by the expansion-specific model across the superior colliculus.

We first tested whether variables used to predict imminent collision in the shallow convolutional neural network are encoded in human superior colliculus activity, and compared them to models using the optical variables τ and η. We fit encoding models^34^ of looming motion to predict fMRI signal acquired as participants (*N* = 15) viewed dynamic visual stimuli used for retinotopic mapping^29^ (see Methods). The visual stimuli included four types of motion: clockwise and counterclockwise sweeping wedges in addition to contracting and expanding rings. These stimuli uniquely activated units in the convolutional network depending on their receptive field (Figure S1). Because expanding rings involve symmetric radial expansion, a hallmark of looming that activates the superior colliculus^26,35^, we hypothesized responses to expanding rings should be best explained by encoding models utilizing features that are useful for detecting imminent collision.

Accordingly, we also compared the performance between two varieties of each encoding model: a stimulus-general version trained to identify mappings between representations of looming and human brain activity using responses to all four stimulus types, and an expansion-specific version trained to identify mappings using only responses to optical expansion. If neural populations in the human superior colliculus responses encode visual looming, then the model trained to predict patterns of BOLD response from optical expansion alone should outperform the stimulus-general model with the same parameters, whereas regions that are sensitive to visual motion more broadly, such as primary visual cortex^36,37^, should be best predicted by the stimulus-general version of the model.

We found that an expansion-specific encoding model built using features from the collision detection model predicted BOLD responses in the superior colliculus (leave-one-subject-out cross-validated *r* = .119, *SE* = .039, 99.0% of noise ceiling, *p* < .001, permutation test; Figure 1D), and that it outperformed its associated stimulus-general model (Δ*r* = .046, *SE* = .021, 63.0% change, *p* = .020, permutation test; responses of individual units are shown in Figure S2). Critically, the enhanced performance of the expansion-specific collision detection encoding model was greater than that of the contraction- and wedge-specific models on matched stimuli (Δ*r* = .092, *SE* = .031, *p* < .001, 158.2% change, permutation test; Figure 1D). Further, adding estimates of looming based on the optical variables τ and η to the encoding model did not improve prediction (Δ*r* = .003, *SE* = .004, 2.7% change, *p* = .198, permutation test, see Supplemental Table 1), demonstrating that these variables do not capture aspects of superior colliculus function beyond those learned by the collision detection model.

Because the superior colliculus receives inputs from primary visual cortex, we next evaluated whether the sensitivity of the superior colliculus to looming is distinct from cortical processing of visual motion. We did so by testing whether representations of looming from the collision detection model differ in their ability to predict responses in superior colliculus and primary visual cortex (V1). This comparison provides a strong analytical control because V1 is sensitive to motion generally, but does not selectively respond to coherent motion. The expansion-specific collision detection encoding model robustly predicted BOLD responses in V1 (*r* = .368, *SE* = .025, 83.0% of noise ceiling, *p* < .001, permutation test), and outperformed its associated stimulus-general model (Δ*r* = .050, *SE* = .010, 15.8% change, *p* < .001, permutation test). We also observed a relative improvement of the looming-specific collision detection model compared to its associated stimulus-general version in V1 compared to the contraction- and wedge-specific models (Δ*r* = .043, 13.1% change, *SE* = .010, *p* = .002, permutation test; Figure S3).

To compare performance between the superior colliculus and primary visual cortex on a balanced scale, as they have different sources of noise and hemodynamics, we estimated the noise ceiling for each region of interest (see Methods) and scaled correlation coefficients separately for each region based on these estimates. Testing on these adjusted values demonstrated that the relative boost in performance of the expansion-specific collision detection model over its stimulus-general version was larger in the superior colliculus than in V1 (*6*1.7% of noise ceiling, *SE* = 26.5%, *p* = .002, permutation test). Taken together, these results show that whereas V1 responses more generally encode information about visual motion, regardless of its coherence and direction, patterns of BOLD activity in the human superior colliculus encode representations of looming motion that has been linked to defensive behavior across species.

### Representations of looming predict defensive blinking in infants

To investigate whether the shallow convolutional network can characterize putatively fear- or threat-related behaviors that depend on superior colliculus function, we evaluated whether it predicts defensive blinking in human infants. Infants develop a propensity to blink in the face of looming stimuli beginning at 4-6 months^23^. Defensive blinking is selective to impending collision and could involve looming computations like the ones modeled by our shallow neural network. If this is the case, and the shallow neural network contains representations of looming that are functionally similar to those used by newborn infants, then infants’ tendency to blink while viewing looming objects should be related to model-estimated collision probability on each frame.

Analyzing defensive blinking in response to visually looming objects (see Figure S4 for timeseries data), we found that collision probability predicted blink count across all frames (*beta* = .427, *SE* = .038, Poisson regression, *p* < .001, permutation test; Figure 2D). To quantify the strength of this relationship, we leveraged the neural network’s stronger activation to faster-approaching stimuli and tested whether infants are similarly sensitive to the velocity of looming stimuli. We found that time points at the end of videos that consistently produced defensive blinking (≥ 5 blinks, see Methods) could be accurately discriminated from other portions of the video (area under the ROC curve (AUROC) = .902, *SE* = .025, *p* < .001, permutation test), and that discriminability increased with object speed (Kendall’s τ = .657, *p* = .046, permutation test; Figure 2E).

**Figure 2.**
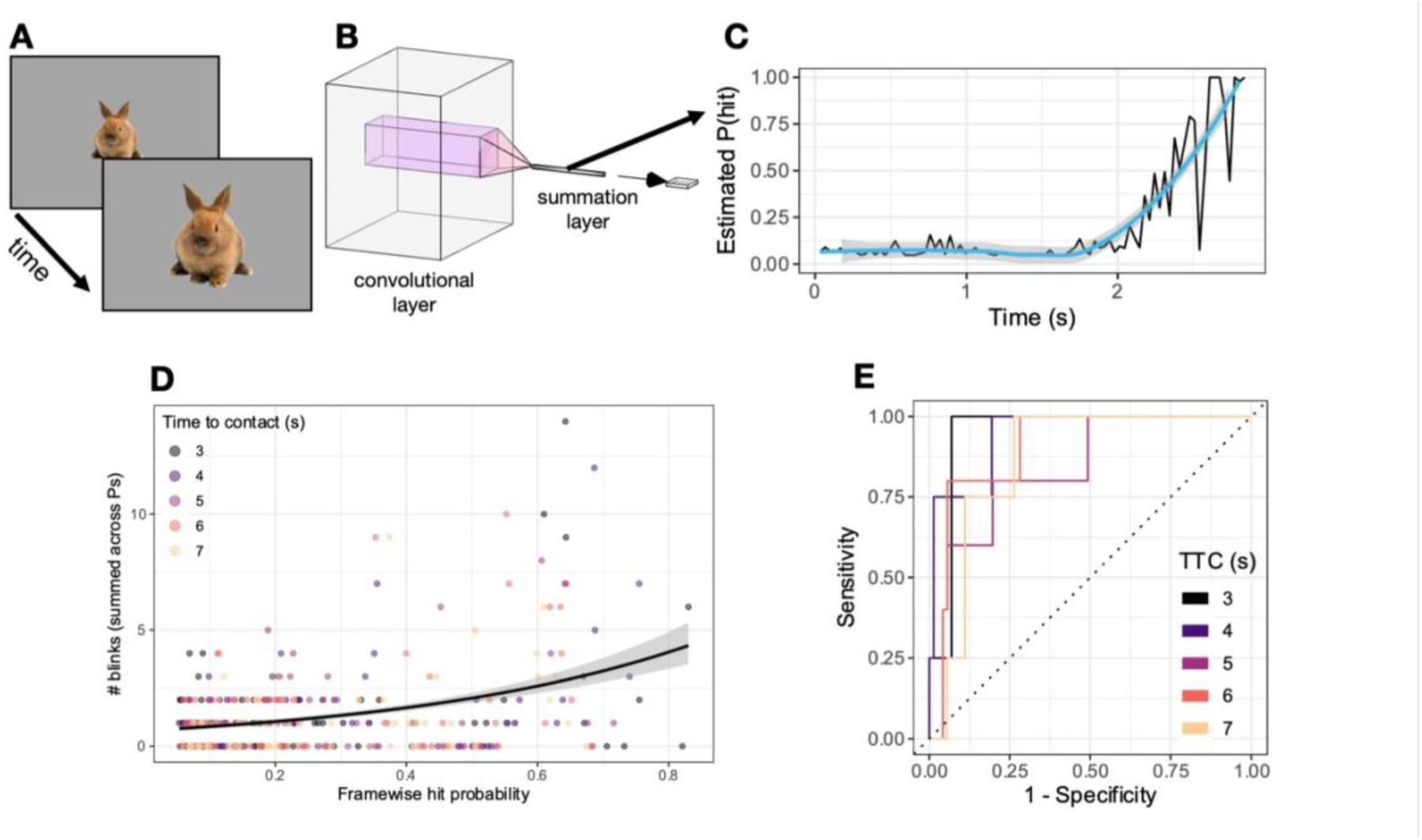
Representations of looming predict defensive blinking to looming objects in infants. **(A)** Videos of looming objects generated by radially expanding static images over time to simulate the appearance of approach motion. **(B)** Depiction of the convolutional neural network, which rectifies and sums unit activations and then applies a softmax activation function to estimate collision probability on each frame. **(C)** Extracted collision probabilities for one representative video. **(D)** Videos with varying apparent times-to-contact showed that greater looming collision probability was associated with increased blinking on a given frame (Poisson regression). **(E**) Receiver operating characteristic curves showing separability of “high-blink” (≥ 5 blinks) and “low-blink” (< 5 blinks) frames

Unlike models of superior colliculus responses, models of defensive blinking based on τ and η were highly predictive (τ: beta = .493, *SE* = .041, *p* < .001; η: beta = .311, *SE* = .044, *p* < .001; permutation test; Table S2). Combining these variables with outputs from the convolutional neural network improved prediction of blink count (ΔAIC = 149; see Table S2 and Figure S5). Together, these observations show that simple computations based on optical expansion are sufficient to predict velocity-sensitive human defensive responses to dynamic looming stimuli, although among candidate models the convolutional neural network alone predicted superior colliculus responses to optical expansion.

### Representations of looming predict subjective emotion elicited by naturalistic videos

Although our findings are consistent with a large literature studying the neural basis of visual looming detection and accompanying defensive behavior across species, it remains unclear how looming contributes to affective experience in humans. Looming is such a strong threat cue that one can readily imagine one’s emotional response to, say, seeing a ball hurtle toward one’s head, even if the ball does not actually make contact. Even though looming is well-established as an aversive and arousing experience^38,39^, we still lack a mechanistic understanding of how looming relates to subjective experience. For example, looming might predominantly inform experience through its relationship with dimensions thought to form the core of affective experience, namely valence and arousal. Alternatively, looming objects may be more specifically related to the experience of fear, because they activate schemas of impending threat (e.g., approaching predators)^42^. To test this hypothesis, we evaluated whether the convolutional network could identify looming motion from a large database of over 2,000 naturalistic videos^30^ and whether activation in the network predicted emotion ratings to the same stimuli.

We trained a partial least squares classifier to discriminate whether 1,315 clips from the database featured an object approaching the camera, using responses to these stimuli from the looming motion model. We tested this classifier on 332 held-out videos from the same database and confirmed that the model predicted human-coded looming above chance (AUROC = .739, chance = 0.5, *SE* = .0007, *p* = .003, permutation test). To test the extent to which visual looming predicts self-reported emotional experience, we then trained a 20-way linear discriminant analysis classifier to identify the consensus emotion category of the same training videos from their looming representations. We found that representations of looming only predicted the top consensus emotion category in the same held-out testing set, though only weakly (16.9%, *SE* = 2.1%, chance = 13.0%, *p* = .010, permutation test; Figure 3D). The AUROC was .538 (chance = .5, *SE* = .024, *p* = .024, permutation test), showing that looming information could discriminate between a subset of emotion classes, but could not fully disentangle the full set of emotions (see Figure S6 for mappings between specific units and different emotion categories).

**Figure 3.**
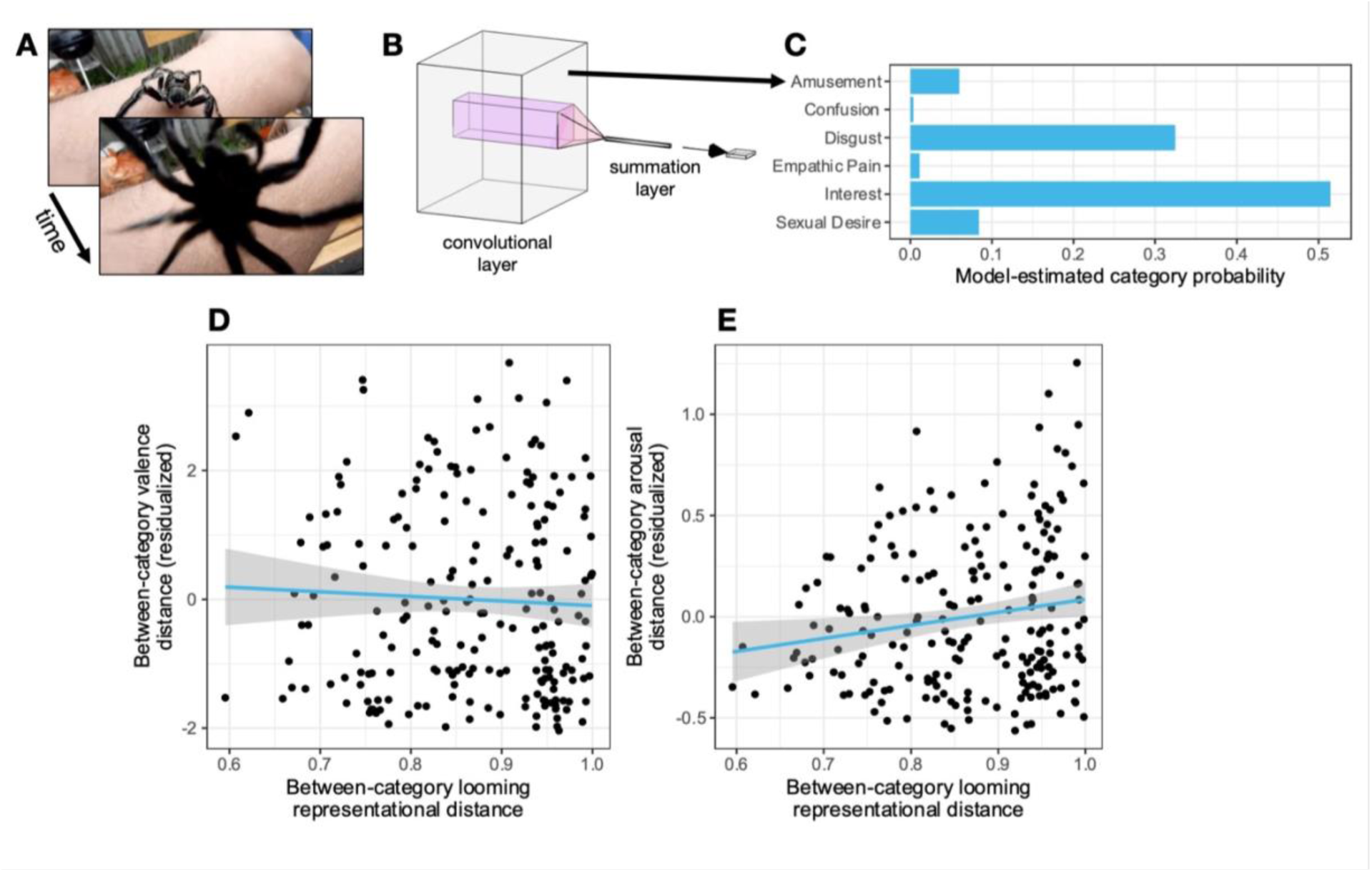
Representations of looming predict subjective emotion evoked by naturalistic videos in adults. **(A)** Participants viewed short video clips depicting a variety of situations. Frames are shown from a stimulus with apparent looming motion. **(B)** We passed the optical flow from these videos through the same convolutional neural network and extracted unit activations from the convolutional layer. **(C)** We trained a 20- way linear discriminant classifier to predict the normative emotion category of each video from its looming activations. **(D)** Distance between emotion categories in the collision detection emotion classifier is unrelated to subjective valence, after adjusting for information from the static feature-based emotion classifier. **(E)** Distance between emotion categories in the collision detection classifier is associated with subjective arousal, after adjusting for information from the static visual feature-based classifier. Distance based on static visual features positively correlated with that of subjective fear, arousal, and valence (Table S2).

To assess which dimensions of experience were predicted by looming, we next quantified the extent to which specific emotion categories (e.g., fear) and more general dimensions such as valence and arousal were the basis for classification. To do so, we compared the similarity of predictions in the 20-way classification (Figure S8) to the similarity of self-report ratings of fear, valence, and arousal (a representational similarity analysis^44^; see Figures S9-10 and Table S3). This analysis revealed that the similarity of emotion categories in the looming-based classifier positively correlated with arousal (partial *r* = .169, *p* = .015, permutation test; Figure 3F) but not subjective fear (partial *r* = .110, *p* = .121, permutation test) or valence (partial *r* = .047, *p* = .495, permutation test; Figure 3F). These findings suggest that in this set of naturalistic videos, representations of looming motion that facilitate the detection of imminent collision discriminate emotional experiences along a dimension of subjective arousal.

It is possible that information about looming motion is unique in its contribution to emotional experience. Motion and static visual features (e.g., texture, shape, color) convey different types of threat-relevant information (e.g., threat imminence versus the source and type of threat^45^) and are processed by distinct neural pathways. A rapidly approaching spider can evoke fear both due to its proximity and appearance^42^. To test whether the shallow convolutional neural network predicts emotion ratings independently from information related to static visual features, we compared the performance of the looming motion-based emotion classifier to a deep network that categorizes emotional situations based on the static content of individual video frames^46^. The ability of the looming classifier and the static feature classifier to classify emotion categories were uncorrelated (Kendall’s τ = -.189, *p* = .122, permutation test). Differences in classification accuracy and comparisons of higher order dimensions (see Figure S7) suggest that some emotion categories (e.g., ‘joy’ and ‘fear’) were better predicted by the presence of looming motion, whereas other categories (e.g., ‘craving’ and ‘desire’) were better predicted by the presence of specific visual features, irrespective of how they move in the environment. These findings suggest that the experience of a looming threat may be aversive due to the presence of co-occurring static properties (that may be integrated with motion), rather than looming being innately aversive on its own.

## Discussion

Here we demonstrate how an incredibly simple network architecture can have broad explanatory power, accounting for different neurobehavioral measures across the lifespan. Recent advances using goal-driven optimization with much more complex architectures (on the order of 10^7^ more parameters) to characterize cortical systems involved in object recognition, speech perception, and language processing^47–49^ have been based on the idea that large, overparameterized models are necessary to explain the human mind. The present findings stand in contrast to this approach, illustrating how a much simpler architecture trained with the right objective function—a computational primitive—characterize multiple aspects of human behavior that are not explained by more complex models of cortical brain systems^46^.

We found that representations of optical expansion from a convolutional neural network for collision detection are encoded in human superior colliculus activity. Although prior related research in humans has demonstrated that the superior colliculus responds more strongly to looming compared to receding motion^26^, it has not examined whether brain activity tracks optical variables that can be used to predict imminent collision. Here we found that representations from a shallow convolutional network predict colliculus responses to optical expansion that are consistent with parametric models based on the optical variables τ and η. Direct readouts of such representations in the superior colliculus could drive defensive behavior in humans, similar to synaptic mechanisms identified in rodents that involve connections with the dorsal periaqueductal grey^50^. Future work is needed to determine if similar circuit-level mechanisms are present in humans, and to characterize how the superior colliculus interacts with cortical and subcortical networks to coordinate defensive behavior^51^.

The present results also contribute to a growing body of evidence implicating the dorsal midbrain in emotional experience^52–54^. Several human neuroimaging studies have revealed that the superior colliculus responds to the aversiveness of visual images^55–58^. It is possible that observations from these studies originate from the same underlying representation of aversiveness. However, our present findings suggest this is not likely the case, as the representations of looming that were encoded in the superior colliculus were largely unrelated to differences in self-reported valence. Given the functional distinction between superficial layers of the colliculus which receive inputs from visual cortex, and deeper layers that contain more specialized loom-sensitive neurons^25^, it is plausible that BOLD responses to static images observed in past studies reflect a subset of neural population activity in the superior colliculus that is not specialized for motion.

In contrast to the typical focus on valence as a building block of emotion, the present work highlights the importance of arousal in explaining emotional behavior. Studies that measure self-reported experience identify hedonic valence as the single dimension that best predicts the semantic structure of emotion ^59^. Experience-sampling suggests that adults organize their emotions primarily using valence ^41^, and developmental studies further show that infants and children first distinguish facial expressions and linguistic concepts using valence^60,61^. We found that computations supporting a species-general behavior predominantly relate to subjective arousal, suggesting that primitive aspects of phenomenal experience may be implemented at the level of the human midbrain^54^. More generally, our findings caution against the assumption that certain stimuli which evoke defensive behaviors produce experiences that resemble prototypical instances of fear in adults, because the computations that underlying these behaviors do not strongly predict subjective valence or fear in a broader array of naturalistic stimuli.

Here we have revealed one way in which human emotions could be based on computations conserved across species. Although we have focused on sensory evaluation, our observations provide a sketch of what understanding emotion might look like from a neurocomputational perspective. Precisely characterizing species-general central emotion states^3^ by modeling how environmental and social affordances shape behavior will likely explain a substantial portion of human emotion. By shifting the focus from a small number of apparently simple, interpretable variables to computationally explicit models that match the complexity of the brain^62^, this approach promises to yield new insights into the origins and nature of emotion.

## Materials & methods

### Implementation of the shallow convolutional neural network

We implemented a shallow neural network model originally built to model the Drosophila LPLC2 pathway and trained to identify whether dynamic stimuli are on a collision course with the viewer ^33^. The network takes in a 4D timecourse of visual motion in each of the 4 cardinal directions. The network has two layers that operate on each frame of the timeseries: one convolutional layer, which, once trained, passes a 12 × 12 px outward motion filter over the visual field to generate a 256-unit representation of looming, and one summation layer, which rectifies, sums, and applies a softmax activation function to estimate looming collision probability for that frame.

For each of the studies described below, we first resized the study’s stimuli to 132 × 132 px to yield 256 convolutional units given the filter size and stride parameters. We then estimated each stimulus’ optical flow using the Farneback algorithm as implemented by OpenCV ^63,64^ and re-cast the optical flow from 2D (positive/negative motion in the x and y directions) to 4D (positive motion in each of the cardinal directions, hereafter referred to as cardinal flow) in accordance with the model.

We then adapted the pre-trained collision detection model from operating on fly-like to human-like vision, instantiating it as a 2D convolutional neural network in PyTorch ^65^ that passes the pre-trained 12 × 12 px outward motion filter over the optical flow from a human-watchable video stimulus, with 11 px stride and 0 px padding, to replicate the unit-to-unit visual field overlap from the original fly-like model. We left the summation layer identical to the original model. Finally, we passed each stimulus’ cardinal flow through the modified collision detection model and extracted representations of looming at various stages of the model to map onto human responses (described further for each study below).

### Study 1: Retinotopic fMRI study

#### Overview

In Study 1, we tested whether looming representations in our model were encoded in human superior colliculus BOLD activity. We leveraged whole-brain fMRI responses to dynamic visual stimuli used for retinotopic mapping to maximize potential looming-related variance in superior colliculus activity. We hypothesized that BOLD responses to visual stimuli would be driven by two types of neural populations: retinotopically organized populations in superficial layers that respond irrespective of motion direction ^66^ and populations in intermediate and deep layers of the colliculus that respond primarily to expanding radial motion ^67^. We tested this hypothesis by fitting multivariate encoding models to predict patterns of colliculus response using the shallow convolutional neural network for collision detection as a feature extractor. If the human superior colliculus contains neural populations that code for visual looming, and they are engaged by the retinotopic videos, then encoding model performance should be the highest on models trained and tested specifically on video stimuli that include optical expansion.

#### Experimental paradigm and stimuli

We used a previously published dataset of retinotopic mapping fMRI scans collected on 15 healthy adult participants ^29^. Participants were scanned while viewing four types of dynamic retinotopy stimuli: clockwise and counterclockwise sweeping wedges, and contracting and expanding rings. The stimuli cycled across the visual field with a period of 32 s, with five repetitions per run, with each run lasting 3 minutes.

#### MRI preprocessing

fMRI data were preprocessed using SPM12 in MATLAB ^68,69^. Images were first realigned to the first image of the series using a six parameter, rigid-body transformation ^70^. The realigned images were then normalized to MNI152 space using a 12-parameter affine transformation followed by nonlinear deformations using a three-dimensional discrete cosine transform basis set, as implemented in SPM ^71,72^. No additional smoothing was applied to the normalized images. Normalized images were subsequently temporally bandpass filtered with cutoff frequencies centered around the stimulus frequency (.667/32 and 2/32 Hz).

#### Measurements

We extracted preprocessed BOLD timeseries from a hand-drawn ROI of the superior colliculus ^56,73^, as well as an ROI of V1 from a multimodal cortical parcellation ^74^ as a positive control.

#### Modeling

We passed sequences of cardinal flow from each retinotopic mapping stimulus through the convolutional layer of the collision detection model. We then convolved the timecourse of units in the shallow convolutional neural network to each of the retinotopy stimuli with the SPM double-gamma hemodynamic response function to generate a multivariate encoding model of looming-related BOLD signal. We applied partial least-squares (PLS) regression, implemented through the mixOmics and tidymodels packages in R ^75–77^, to map our looming-predicted BOLD onto observed multivariate BOLD from each ROI separately. We trained the PLS multivariate encoding model on data from 14 participants and then assessed model fit as the Pearson correlation between PLS-predicted BOLD and observed BOLD in the last held-out participant. We cross-validated model fit in a leave-one-subject-out manner by repeating this process for every participant and averaging across repetitions ^78^.

Because the collision detection model contains units that tile the visual field, the resulting BOLD encoding model encodes both retinotopic responses and responses to looming motion. Accordingly, to test for looming specificity, we compared performance between two types of encoding models: a stimulus-general model, with the PLS mapping trained on data from all four stimulus types, and stimulus-specific models, with the PLS mapping trained separately on data from each stimulus type. We expected the stimulus-specific model trained on expanding ring motion would predict superior colliculus responses more so than other stimulus-specific models, or the stimulus-general model.

In order to clarify the nature of the looming representations in our BOLD encoding model, we also compared performance between the neural network encoding model and encoding models predicting BOLD responses as a function of the optical looming variables τ and η. For the retinotopic ring stimuli, we calculated timecourses using τ and η based on the visual angle parameters at which the videos were presented to participants, using formulas from ^79^. We fit this optical variable encoding model, along with several variations using different combinations of predictors (Table S1), using the same method described above.

In order to facilitate comparisons of model performance between the superior colliculus and V1, we adjusted model fit correlations by the noise ceilings from their respective ROIs. In each ROI, we estimated the noise ceiling on each cross-validation fold by calculating the Pearson correlation between the average timeseries of that fold’s training participants and the held-out participant, and averaging across folds. We estimated a separate noise ceiling for each retinotopic stimulation condition and used the highest noise ceiling to normalize all encoding model fit estimates.

We generated block permutation distributions against which to compare the model fit correlations by randomizing TRs of observed BOLD within each stimulus cycle to preserve the autocorrelation structure of the data ^80^. We then re-estimated each shuffled model fit correlation over 5,000 iterations to generate p-values for inference.

### Study 2: Infant behavioral study

#### Overview

In Study 2, we tested whether looming representations in our model could predict infant defensive blinking in response to looming stimuli.

#### Participants

A total of 62 healthy infants participated in this study. Of the 62 infants, four infants looked less than 35% of the (total) trial durations and, thus, were excluded from subsequent analyses. An additional 12 infants failed to complete the study due to fussiness or technical difficulties, leaving 58 infants in the final sample (*range* = 6.2-11.7 months, *M* = 8.7 months; 22 boys and 36 girls). Parents provided written informed consent on behalf of their infants. All procedures were approved by the Institutional Review Board at Emory University.

#### Procedure and stimuli

Infants were tested individually in a dimly lit, soundproof room. Each infant sat in a highchair or on his/her parent’s lap at a distance of approximately 60 cm from a large projection screen (92.5 × 67.5 cm). Parents were instructed to keep their eyes closed and to refrain from interacting with their infants during the study, except for soothing them if they became fussy. Stimuli were videos of a looming two-dimensional image, which were rear-projected onto the screen at eye-level to the infant. Each infant’s face was recorded for later coding using a concealed camcorder placed just under the projection screen. Video feed was transmitted directly to a computer in an adjoining room where an experimenter monitored the session remotely.

Images in each of the videos were of individual animals (snakes, spiders, butterflies, and rabbits; two of each type). Images were selected from an Internet search for their high quality and to match roughly in color and brightness. Images were cropped, resized, and presented against a uniform gray background using Adobe Photoshop CS5 ^81^. Looming videos were created in MATLAB by manipulating the rate of expansion of the image size.

Each trial was experimenter controlled, beginning with a centrally presented attention-getter (e.g., swirling star; randomly selected across trials) that played until infants oriented to the screen. A looming video immediately followed. Each video began with two-dimensional image that expanded to a maximum size of 75° × 59° (visual angle). There was a 1 s inter-trial interval (ITI) consisting of a gray screen.

Videos were created such that the virtual animal approached the infant at one of six velocities, indicating times-to-contact of 3, 4, 5, 6, 7, or 8 s. Velocity was negatively correlated with approach time, such that as approach time increased, the velocity of the virtual object decreased. Infants were presented with a total of 48 trials (randomized).

#### Video coding

High quality videos of each infant were saved digitally. Video frames were coded at 33.33 ms intervals by observers blind to the stimuli presented to infants. All videos were coded by one observer for blinks (and total looking time) on each trial. Eye closures were counted as blinks if the lids of the opened eyes covered at least half of the exposed eye surface ^82^. Incomplete eye closures associated with large head turns were not counted as blinks. Also not counted as blinks were eye closures associated with yawns, sneezes, coughs, and hand movements to or near the face or mouth. A second observer coded a random sample of videos (20%) to assess reliability. Inter-observer reliability was high for the coding of both blinks and looking times (*r*s > 0.9).

#### Measurements

For each looming video stimulus presented to the infants, we summed the total number of blinks made by all infants on each coded frame to generate one timecourse of blink counts per video stimulus. We then further summed the blink count timecourses for each video of a given time-to-contact duration to generate one timecourse of total blink counts per time-to-contact condition (Figure S4).

#### Modeling

We extracted the cardinal optical flow for each looming video stimulus at a frame rate of 33.33 ms/frame, and then passed the flow videos through the convolutional and summation layers of the collision detection model to generate a 1D timecourse of estimated collision probability for each stimulus. We then averaged the timecourses for each video of a given time-to-contact duration to generate one timecourse of looming collision probability per time-to-contact duration.

Then, we used Poisson regression to predict framewise blink counts as a function of framewise collision probability and condition-wise time-to-contact. We generated a permutation distribution against which to compare the coefficient for collision probability by randomizing blink counts across all trials. We then re-fit the Poisson regression and extracted the shuffled coefficient over 10,000 iterations to generate *p*-values for inference.

Similar to Study 1, we compared this Poisson model to another Poisson model with the optical variables τ and η added as predictors, in order to clarify the nature of the looming representations encoded in collision probability. First, we estimated timecourses of τ and η for each stimulus video, based on the visual angle parameters at which the videos were presented to participants. We then included these timecourses as predictors in an expanded Poisson model. We ran this model as a principal components regression, applying PCA to the three collision variables (collision probability, τ, and η) and including the three rotated components as predictors in the Poisson regression along with condition-wise time-to-contact.

Finally, we examined the potentially threshold-like relationship between blink counts and collision probability by using collision probability to classify frames as “high-blink” (5 or more blinks across infants/stimuli on that frame, to isolate trials where blinks were most likely to be defensive) or “low-blink” (fewer than 5 blinks). We calculated the area under the receiver operating curve (AUROC) both overall and as a function of time-to-contact condition, using tools implemented in the tidymodels family of R packages ^76^. We evaluated whether AUROC varied with time to collision by calculating Kendall’s τ between the observed rank-ordering of times-to-contact based on AUROC (highest to lowest) and duration (3 s to 7 s). We generated a non-parametric sampling distribution for overall AUROC by bootstrap resampling and re-calculating AUROC over 10,000 iterations. We also generated a permuted distribution against which to compare the observed AUROC by randomizing binarized blink counts across all trials and re-estimating AUROC over 10,000 iterations. Similarly, we generated a block permutation distribution against which to compare the observed Kendall rank correlation between time-to-contact and AUROC by randomizing binarized blink count within each time-to-contact condition. We then re-estimated the shuffled AUROC for each time-to-contact and re-calculated Kendall’s τ over 10,000 iterations to generate *p*-values for inference.

### Study 3: Adult behavioral study

#### Overview

In Study 3, we tested whether looming representations in our model could predict normative self-report affect ratings in response to short, naturalistic videos.

#### Stimuli and behavioral measurements

We used a previously published subset of short, naturalistic videos and normative emotion ratings ^30^. Each video was rated by approximately 10 raters (range = [9, 17]), each of whom reported the categorical emotions elicited by the video, as well as 9-point valence and arousal ratings. For each video, we took its most frequently selected categorical emotion label, and its mean valence and arousal ratings. Videos spanned 20 consensus emotion categories. To quantify ground-truth looming, author PAK coded each video for the presence of objects approaching the camera.

#### Modeling

We resampled each video stimulus to a standard frame rate of 10 fps and passed the cardinal flow from each video stimulus through the convolutional layer of the collision detection model to yield 256 timecourses of activations per video. Next, we flattened each video’s looming representation along the time dimension. The original looming model tends to increase activation over time for “hit” stimuli as the stimuli approach the viewer and activate an increasing number of units across the visual field. Accordingly, we assumed that stronger looming activations would have a more positive slope over time. We calculated the linear slope of each unit’s timecourse over time, generating a looming representation of 256 unit activation slopes per video. We applied partial least squares classification, implemented through the mixOmics and tidymodels packages in R, to classify whether each video was coded as containing looming motion using its 256 looming activation slopes. We trained the partial least squares classifier using a prior training split of 1,315 videos ^46^. We then applied linear discriminant analysis, implemented through the MASS and tidymodels packages in R, to classify each video’s consensus emotion category (out of 20) using its 256 looming activation slopes. We trained the linear discriminant classifier using the same prior training split as the partial least squares looming classifier. All model performance statistics are reported as evaluated on the associated prior held-out testing split of 332 videos.

We compared the emotion classification performance of the looming model to the performance of a deep convolutional neural network originally trained to classify stimulus-elicited emotions based on their static image features ^46^. Because that model was originally used to identify the emotion categories of individual video frames, we calculated video-wise category predictions by averaging each of the 20 emotion class probabilities across each frame of the video and taking the emotion category with the highest across-video average probability. We generated non-parametric sampling distributions for our statistics by bootstrapping and re-calculating classification accuracy, over 10,000 iterations. We also generated non-parametric null distributions against which to compare classification accuracies by permuting the consensus emotion category labels across videos and re-calculating shuffled classification accuracy, over 10,000 iterations. Finally, we generated a permutation distribution against which to compare Kendall’s τ for category rankings by model AUROC by randomizing consensus emotion category label the across videos. We then re-estimated shuffled category-specific AUROCs for both the looming model and the static image model and re-calculated a shuffled Kendall’s τ over 10,000 iterations to generate *p*-values for inference.

We used representational similarity analysis ^44^ to assess whether the representations learned by the emotion classification models encoded information consistent with valence and/or arousal. For both the looming motion-based and static visual feature-based classifiers, we calculated the representational distance between every pair of emotion categories. For a given emotion classification model and pair of emotion categories, we calculated the distance as 1 minus the average pairwise Pearson correlation between the 20 class probabilities for any two videos from those two emotion categories. We then used linear regression to predict between-category distances in mean valence ratings from distances from both convolutional networks, allowing us to assess the independent contributions of information gleaned from optical flow and static visual features. From this regression, we estimated the partial correlation coefficients that identify the relationship between representations of looming and valence (accounting for static visual features), and between representations of static visual features and valence (accounting for looming). We conducted similar regressions using mean ratings of arousal and fear, and extracted partial correlation coefficients using the same approach. We generated permutation distributions against which to compare these partial correlation coefficients ^83,84^, calculating randomized partial correlation coefficients over 10,000 iterations to generate p-values for inference.

## Acknowledgments

We thank Baohua Zhou for assistance with configuring the shallow neural network model, and the ECCO Lab at Emory University for helpful feedback on the project. This work was supported by the National Institutes of Health Institutional Research and Career Development Award (IRACDA) grant K12GM000680 to MKT.

## Additional information

### Author contributions

Conceptualization: PAK, MKT Methodology: PAK, SFL, MKT Investigation: VA

Formal analysis: PAK, MKT Software: MKT Visualization: MKT

Project administration: PAK, SFL Supervision: PAK, SFL

Writing – original draft: PAK, MKT

Writing – review & editing: VA, PAK, SFL, MKT

### Data and materials availability

All data and materials that were generated for this study will be posted on Open Science Framework and all code will be posted on GitHub by time of publication.

