## Supplemental Figures and Tables for "Visual looming is a primitive for human emotion"

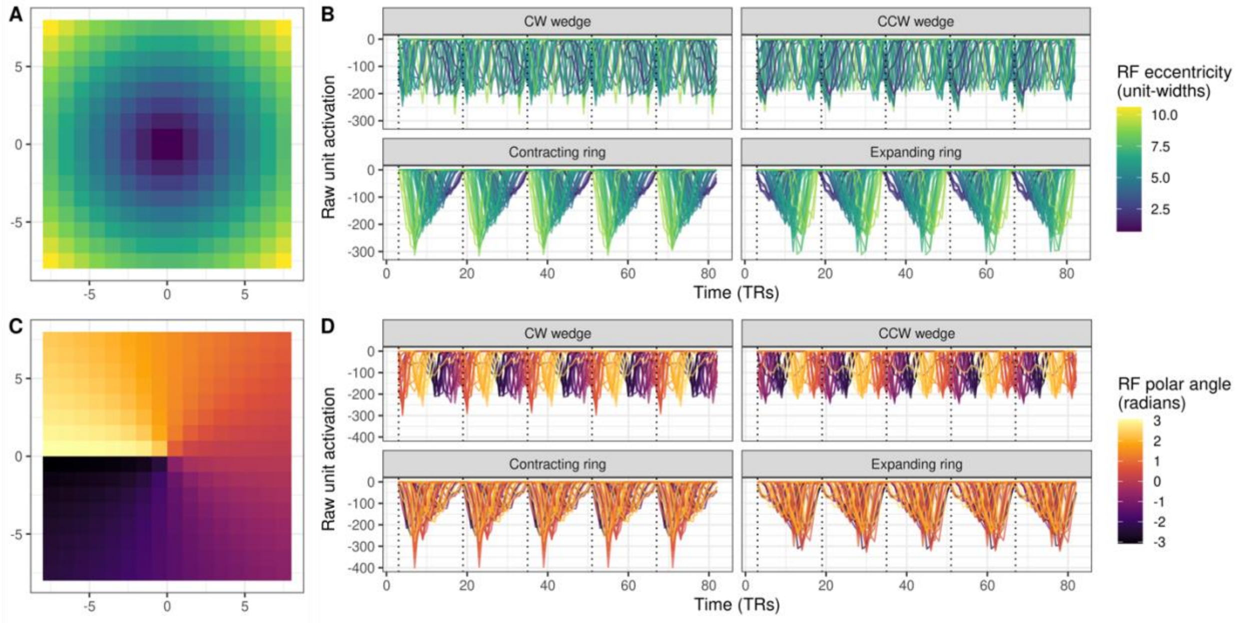


**Supplementary Figure 1.** Activation profiles in the collision detection fMRI encoding model as a function of unit location in the visual field. **(A)** Mapping of units based on their eccentricity. **(B)** During periods of retinotopic ring stimulation (bottom panels), units show rolling phasic activation in order of eccentricity. **(C)** Mapping of units based on polar angle. **(D)** During periods of sweeping wedge stimulation (top panels), units show rolling phasic activation in order of polar angle.

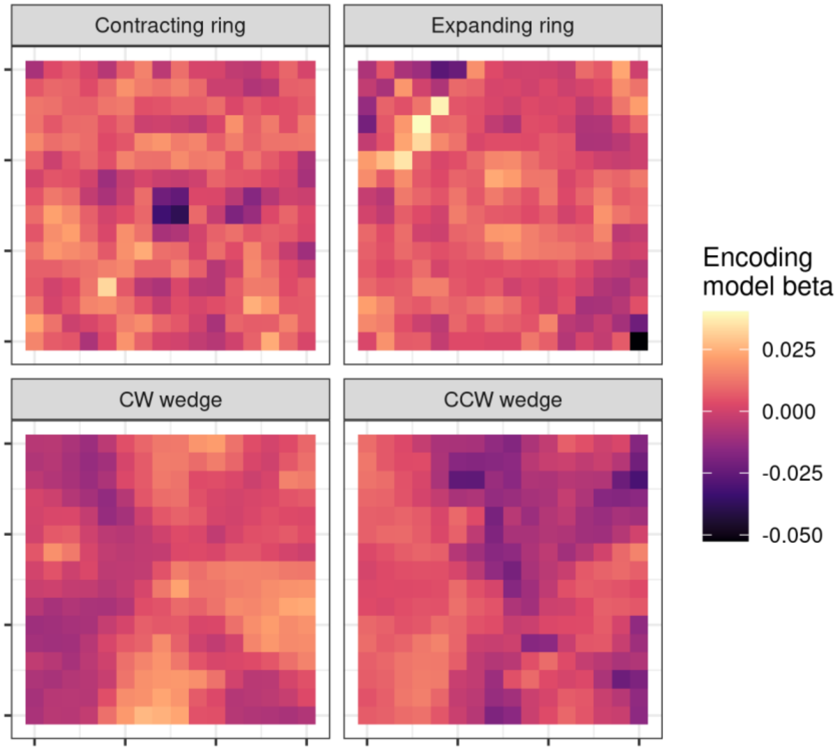


**Supplementary Figure 2.** Structure coefficients from stimulus-specific superior colliculus encoding models. Each heatmap displays group average PLS betas that map activity in the convolutional neural network to the average superior colliculus BOLD response. Units are organized based on their "receptive field" location in the visual field. Warm colors indicate areas of the visual field where greater looming-related model activation tracks positively with superior colliculus activity.

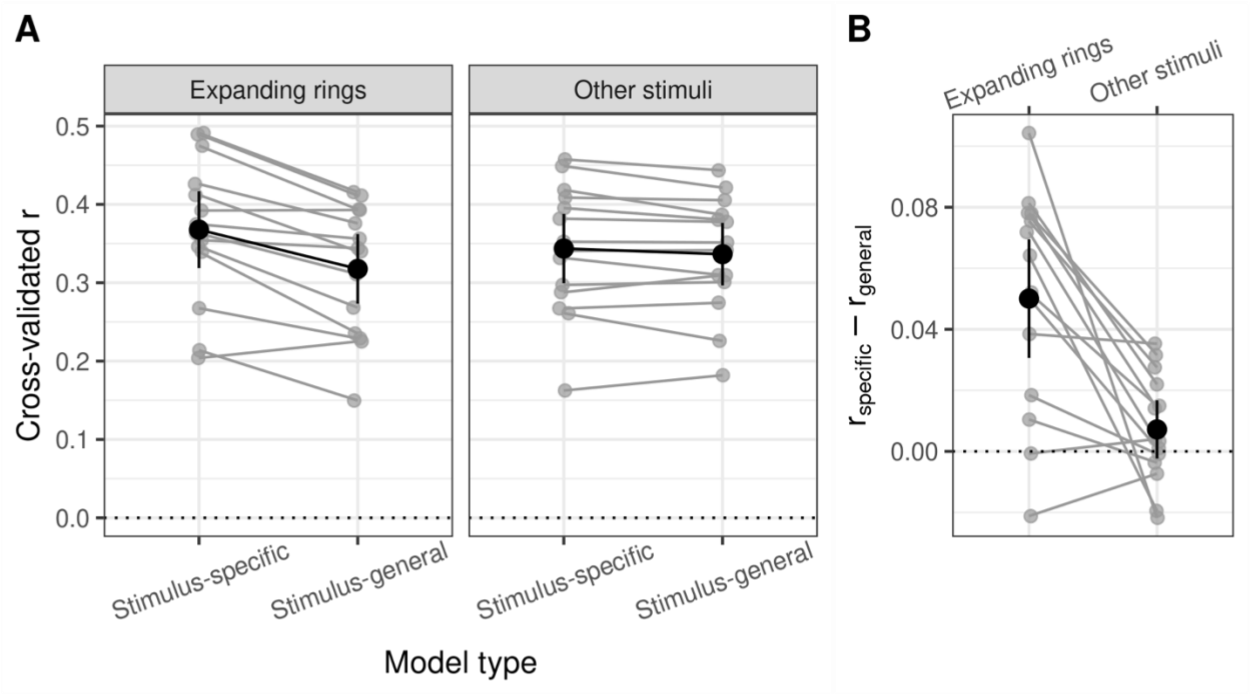


**Supplementary Figure 3.** Collision detection encoding model performance in primary visual cortex. **(A)** Model performance using leave-one-subject-out cross-validated Pearson's r between encoding model-predicted and observed BOLD. Gray points and lines show model fit estimates for each held-out subject. Black summary points and error bars show mean ±2 standard errors across cross-validation folds. The expansion-specific model of visual cortex activity outperforms a stimulus-general model on the same data (left subplot). **(B)** Difference in model fit between the stimulus-specific and stimulus-general encoding models for expanding rings.

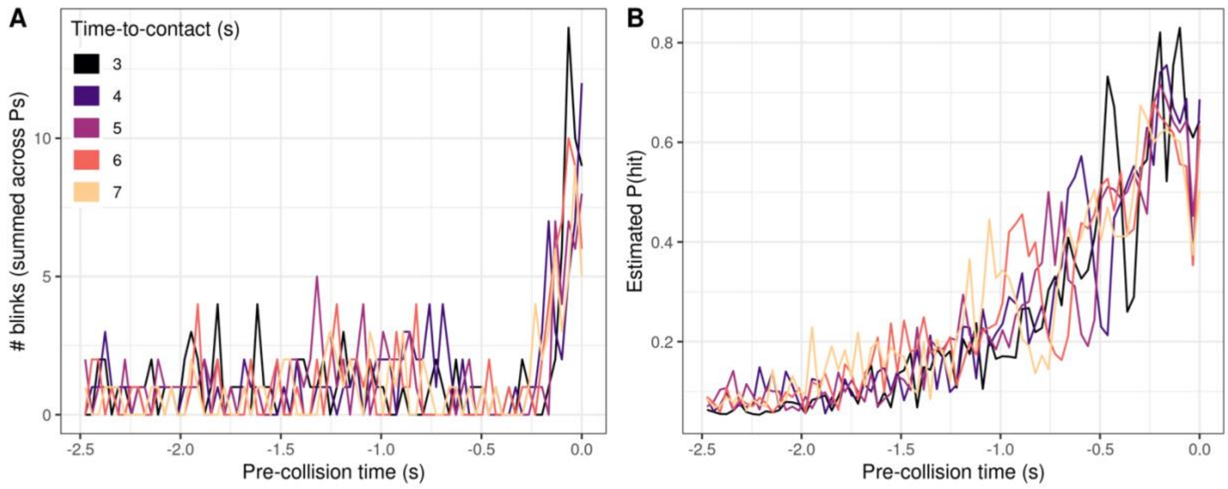


**Supplementary Figure 4.** Infant defensive blinking in response to looming objects. **(A)** Observed infant blink counts and **(B)** model-estimated collision probabilities over time in response to looming objects as a function of apparent time-to-contact. Data for the last 75 frames of each stimulus are shown, time-locked to the point of apparent collision at end of each looming video. “High-blink” trials (>=5 blinks) are clearly concentrated at the end of the stimuli, when impending collision is nearest.

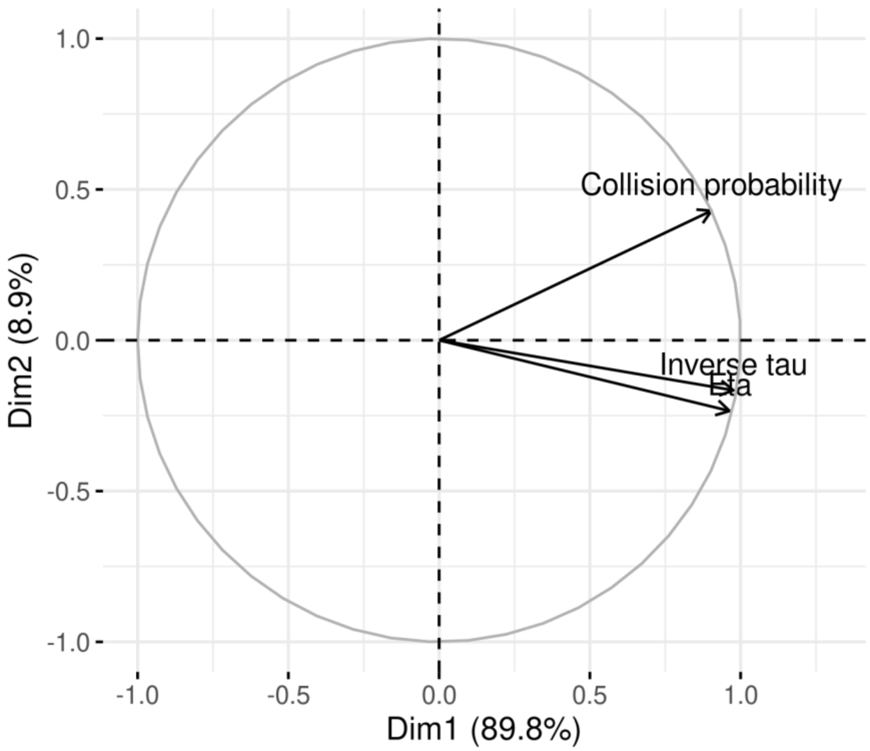


**Supplementary Figure 5.** Correlation circle plot from a principal component analysis on framewise collision probability from the neural network model and optical variables $\tau$ and $\eta$, all estimated on videos of artificially looming objects. The first principal component, shown on the x-axis of the unit loading circle, accounts for 89.8% of the total covariance. All three quantities project strongly onto the first principal component, but collision probability also projects onto the second principal component, shown on the y-axis.

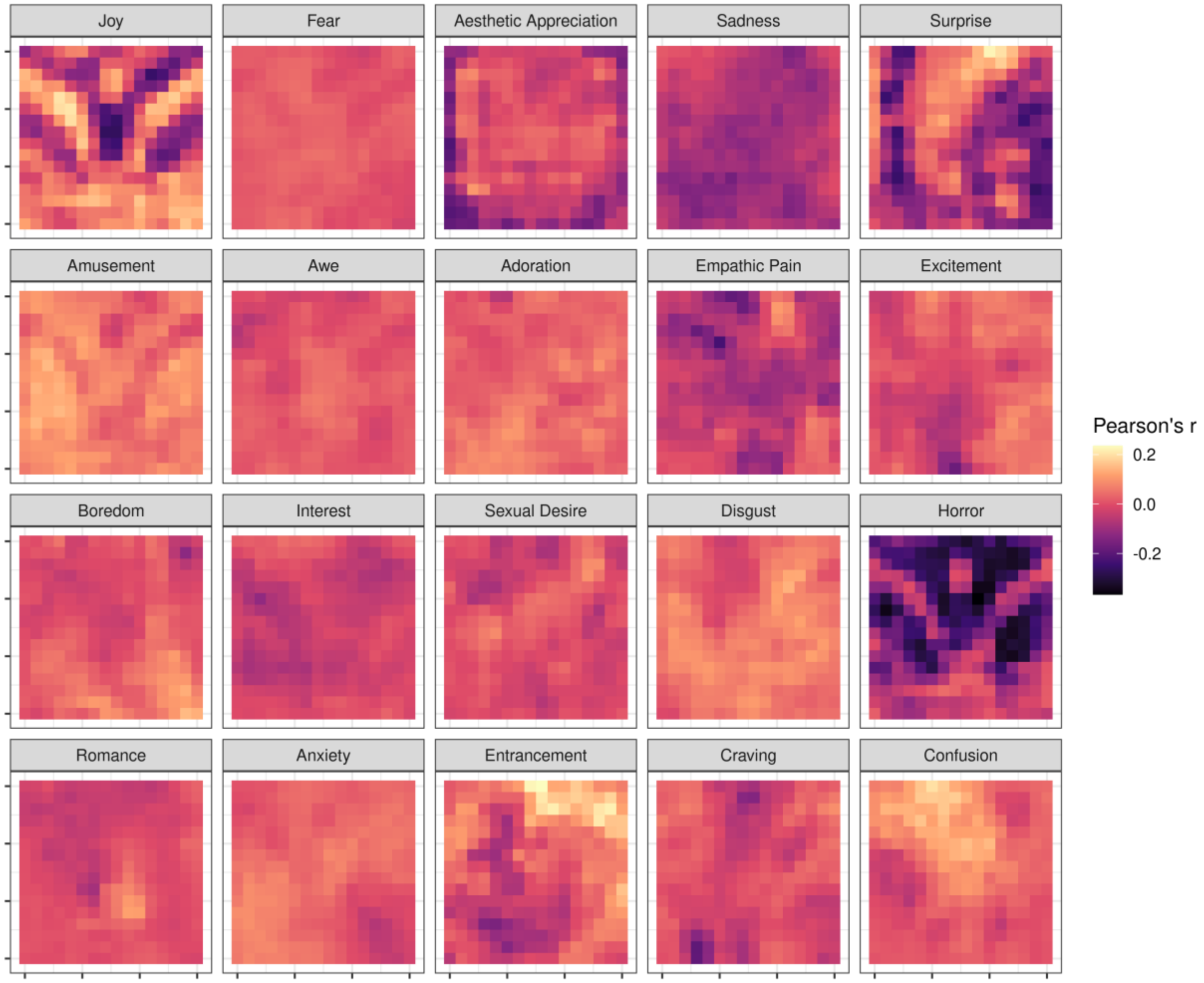


**Supplementary Figure 6.**Structure coefficients of the looming motion-based 20-way emotion classifier. Each emotion category heatmap cell displays the correlation (Pearson's *r*) between each neural network unit activation slope and the subsequent probability of classifying a video as that emotion, across videos. Units are organized based on their "receptive field" location in the visual field, to show spatial patterns of outward motion that characterize various emotion categories. Panels are ordered by the one-vs-all AUROC for each emotion category from highest to lowest discriminability (see Supplementary Figure 7). "Hotter" areas are areas of the visual field where greater looming-related model activation tracks positively with the probability of classifying a video into that emotion category.

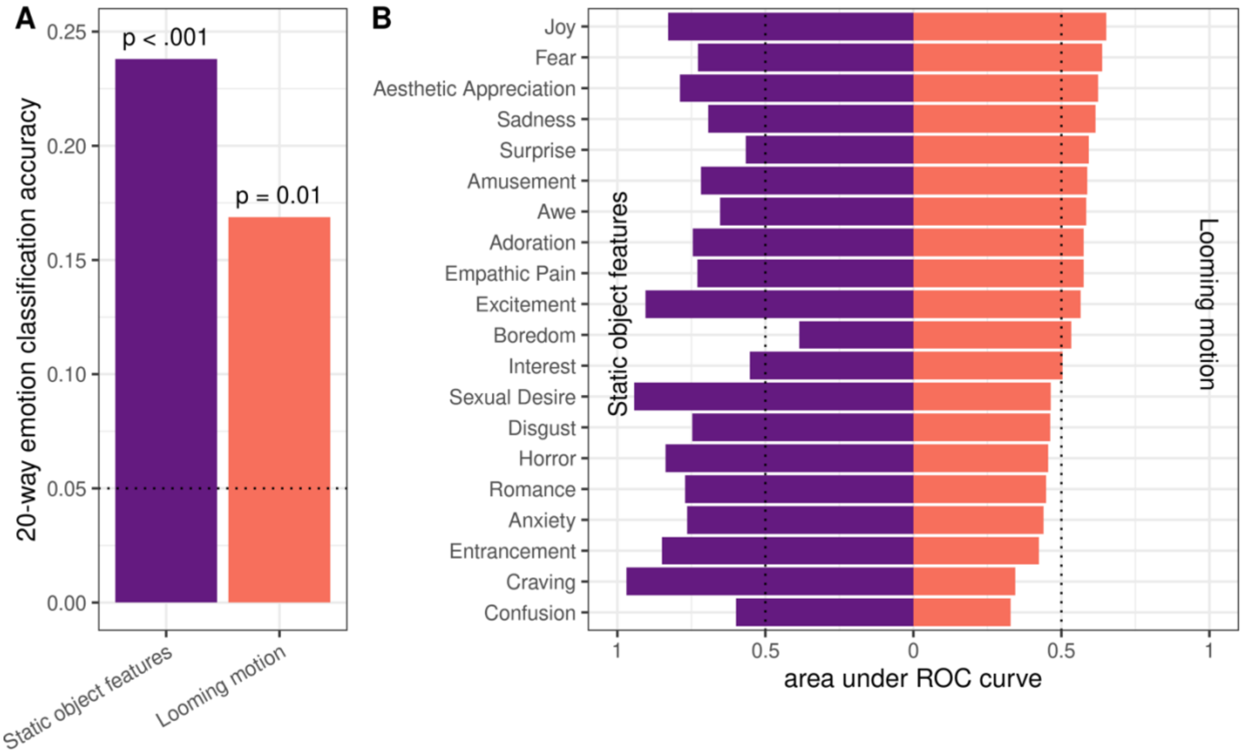


**Supplementary Figure 7.** Performance of the looming-motion-based and static-feature-based emotion classifiers. **(A)** The two models both classify videos into consensus emotion categories above chance (0.05, dotted reference line), but the shallow looming-motion-based classifier (orange, right) does not perform as well as the deep static-feature-based classifier (purple, left). **(B)** The looming-motion-based (orange, right) and static-feature-based (purple, left) classifiers show different patterns of discriminability for specific emotion categories. For example, the looming classifier was relatively better at discriminating videos labeled as "fear" (looming classifier: AUROC = .638, SE = .066, 2nd out of 20; static classifier: AUROC = .727, SE = .064, 13th out of 20) and "surprise" (looming classifier: AUROC = .593, SE = .106, 5th out of 20; static classifier: AUROC = .565, SE = .137, 18th out of 20). Meanwhile, the static classifier was relatively better at discriminating videos labeled as "craving" (looming classifier: AUROC = .345, SE = .091, 19th out of 20; static classifier: AUROC = .969, SE = .017, 1st out of 20) and "sexual desire" (looming classifier: AUROC = .465, SE = .074, 13th out of 20; static classifier: AUROC = .943, SE = .018, 2nd out of 20). Area under the receiver operating characteristic curve (AUROC) was estimated separately for each emotion category in a one-vs-all scheme.

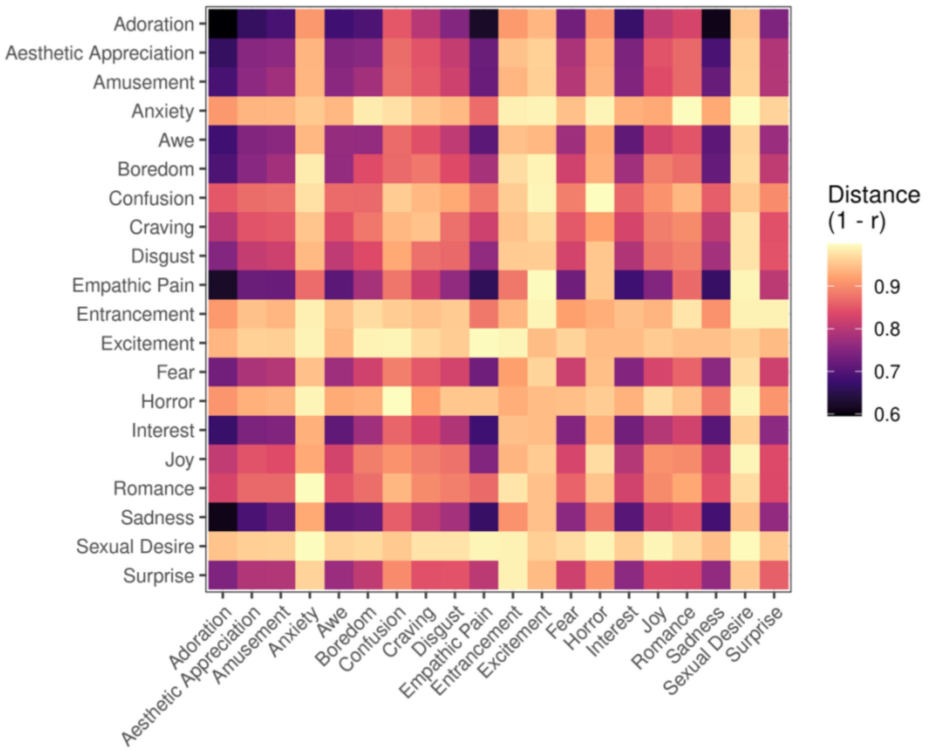


**Supplementary Figure 8.** Distances between emotion categories in the looming-motion-based emotion classifier. Heatmap displays the average correlation distance (1 - Pearson's $r$) based on model-predicted class probabilities between each pair of emotion categories.

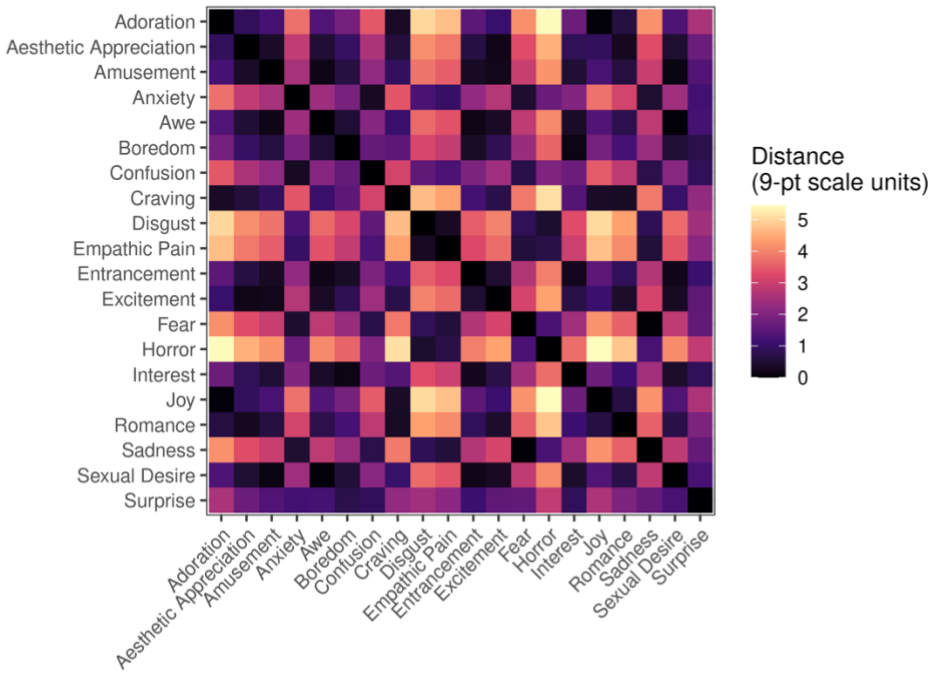


**Supplementary Figure 9.** Distances between emotion categories based on self-reported valence. Heatmap displays the difference in average valence ratings (on a 9-point scale) for each pair of emotion categories.

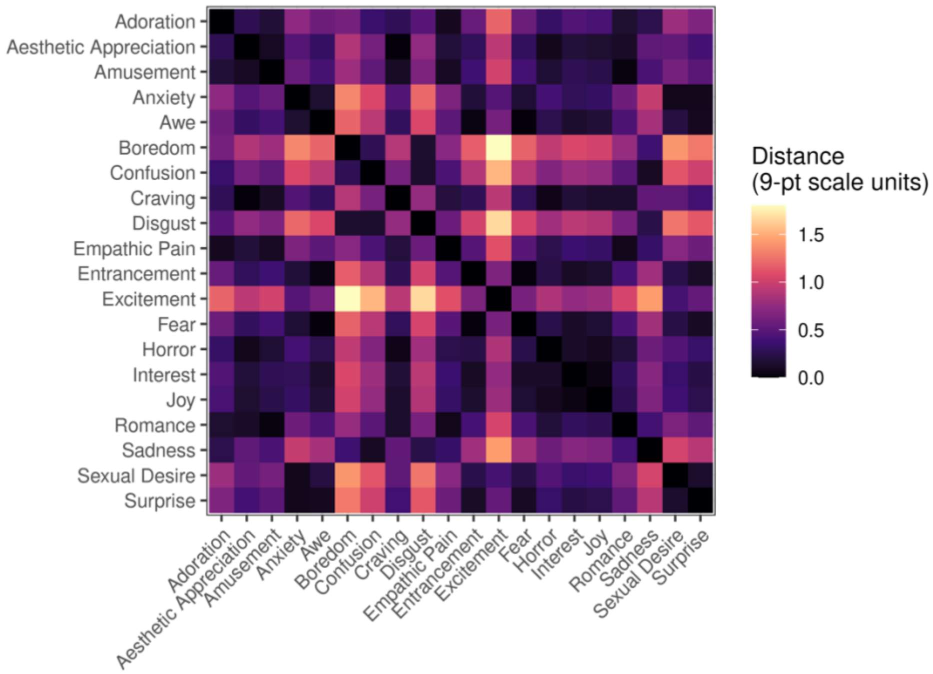


**Supplementary Figure 10.** Distances between emotion categories based on self-reported arousal. Heatmap displays the difference in average arousal ratings (on a 9-point scale) for each pair of emotion categories.

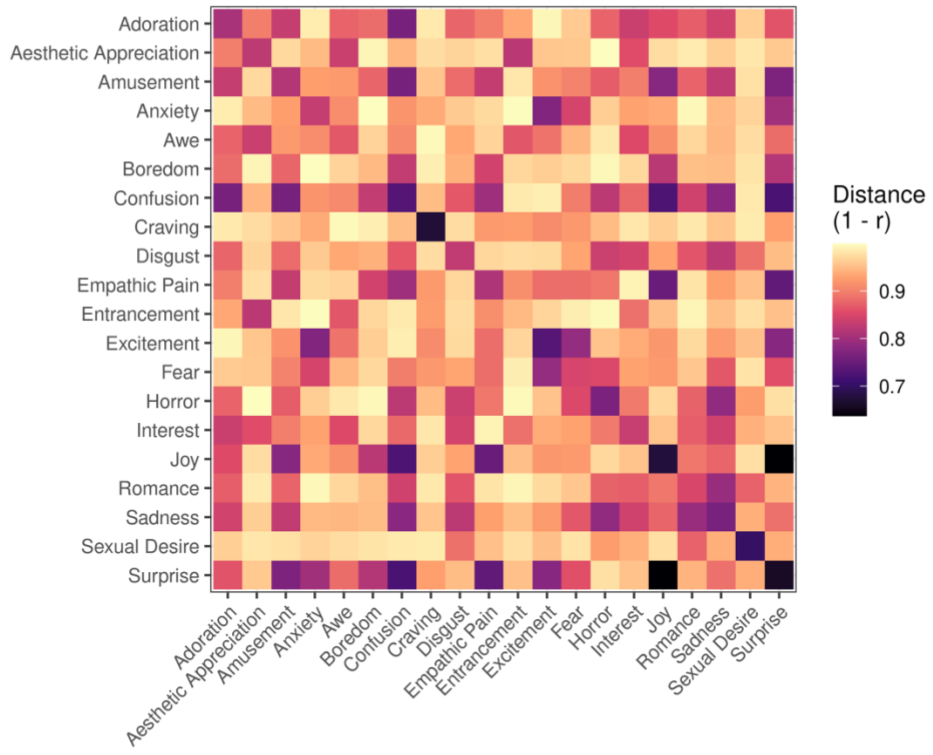


**Supplementary Figure 11.** Distance between emotion categories in the static-feature-based emotion classifier. Heatmap displays the average correlation distance (1 - Pearson's *r*) based on model-predicted class probabilities between each pair of emotion categories.

| ROI | Model predictors | Stimulus condition | Model type | Cross-validated *r* | Standard error |
|---|---|---|---|---|---|
| SC | Tau and Eta | Expanding rings | Stimulus-specific | 0.117 | 0.0516 |
| SC | Tau and Eta | Expanding rings | Stimulus-general | -0.0267 | 0.0192 |
| SC | Eta | Expanding rings | Stimulus-specific | -0.00224 | 0.0157 |
| SC | Eta | Expanding rings | Stimulus-general | -0.018 | 0.012 |
| SC | CNN Activation | Expanding rings | Stimulus-specific | 0.119 | 0.0385 |
| SC | CNN Activation | Expanding rings | Stimulus-general | 0.0733 | 0.0345 |
| SC | CNN Activation | Other stimuli | Stimulus-specific | -0.00286 | 0.0183 |
| SC | CNN Activation | Other stimuli | Stimulus-general | 0.0427 | 0.0204 |
| SC | CNN Activation, Tau, and Eta | Expanding rings | Stimulus-specific | 0.123 | 0.0381 |
| SC | CNN Activation, Tau, and Eta | Expanding rings | Stimulus-general | 0.0731 | 0.0325 |
| SC | CNN Activation, Tau, and Eta | Other stimuli | Stimulus-general | 0.0309 | 0.0209 |
| SC | CNN Activation + Eta | Expanding rings | Stimulus-specific | 0.123 | 0.0382 |
| SC | CNN Activation + Eta | Expanding rings | Stimulus-general | 0.0718 | 0.0322 |
| SC | CNN Activation + Eta | Other stimuli | Stimulus-general | 0.0394 | 0.0205 |
| SC | CNN Activation + Tau | Expanding rings | Stimulus-specific | 0.123 | 0.0381 |
| SC | CNN Activation + Tau | Expanding rings | Stimulus-general | 0.0721 | 0.0315 |
| SC | CNN Activation + Tau | Other stimuli | Stimulus-general | 0.0374 | 0.0205 |
| SC | Tau | Expanding rings | Stimulus-specific | -0.0083 | 0.0126 |
| SC | Tau | Expanding rings | Stimulus-general | 0.00901 | 0.0317 |
| V1 | Tau and Eta | Expanding rings | Stimulus-specific | 0.195 | 0.0137 |
| V1 | Tau and Eta | Expanding rings | Stimulus-general | 0.0713 | 0.0162 |
| V1 | Eta | Expanding rings | Stimulus-specific | 0.124 | 0.00911 |
| V1 | Eta | Expanding rings | Stimulus-general | 0.0104 | 0.016 |
| V1 | CNN Activation | Expanding rings | Stimulus-specific | 0.368 | 0.0245 |
| V1 | CNN Activation | Expanding rings | Stimulus-general | 0.318 | 0.0222 |
| V1 | CNN Activation | Other stimuli | Stimulus-specific | 0.344 | 0.0145 |
| V1 | CNN Activation | Other stimuli | Stimulus-general | 0.336 | 0.0127 |
| V1 | CNN Activation, Tau, and Eta | Expanding rings | Stimulus-specific | 0.366 | 0.0253 |
| V1 | CNN Activation, Tau, and Eta | Expanding rings | Stimulus-general | 0.326 | 0.0231 |

| | | | | | |
|---|---|---|---|---|---|
| V1 | CNN Activation, Tau, and Eta | Other stimuli | Stimulus-general | 0.335 | 0.0129 |
| V1 | CNN Activation + Eta | Expanding rings | Stimulus-specific | 0.366 | 0.0253 |
| V1 | CNN Activation + Eta | Expanding rings | Stimulus-general | 0.32 | 0.0223 |
| V1 | CNN Activation + Eta | Other stimuli | Stimulus-general | 0.335 | 0.0129 |
| V1 | CNN Activation + Tau | Expanding rings | Stimulus-specific | 0.366 | 0.0253 |
| V1 | CNN Activation + Tau | Expanding rings | Stimulus-general | 0.326 | 0.0233 |
| V1 | CNN Activation + Tau | Other stimuli | Stimulus-general | 0.335 | 0.0129 |
| V1 | Tau | Expanding rings | Stimulus-specific | 0.124 | 0.00926 |
| V1 | Tau | Expanding rings | Stimulus-general | 0.102 | 0.0166 |

**Supplementary Table 1.** Cross-validated performance (Pearson's *r* between predicted and actual BOLD timecourses) for all partial least squares encoding models. Table columns are organized by ROI (superior colliculus and V1), included model predictors (models with one or both optical variables, the primary model with neural network activations, and combined models with neural network activations and optical variables).

| Model predictors | AIC |
|---|---|
| CNN Activation, Tau, and Eta | 1113 |
| Tau | 1238 |
| CNN Activation and Tau | 1239 |
| CNN Activation and Eta | 1253 |
| CNN Activation | 1262 |
| Eta | 1329 |

**Supplementary Table 2.** Model performance (AIC) for all models predicting infant blink count as a function of $\tau$ and $\eta$ and/or neural network collision probability. Rows are ordered by AIC from lowest (best model performance) to highest. Coefficients from models in rows 2, 5, and 6 (the models with only one predictor of interest) are reported in the main text.

| Model type | Outcome | Partial r | p-value |
|---|---|---|---|
| Looming motion | Arousal | 0.169 | 0.0153 |
| Looming motion | Valence | 0.0477 | 0.495 |
| Looming motion | Fear | 0.11 | 0.121 |
| Static visual features | Arousal | 0.237 | .0008 |
| Static visual features | Valence | 0.174 | 0.0115 |
| Static visual features | Fear | 0.184 | 0.0079 |

**Supplementary Table 3.** Representational similarity analysis results relating subjective arousal, valence, and fear to visual model representations of 20 emotion categories (as depicted in Figures S7-10). Rows 1-3 for the looming-motion-based classifier are also reported in the main text. The table shows further that representational similarity based on static visual features positively correlated with that of subjective fear, arousal, and valence (rows 4-6). Partial r is reported for each classifier & outcome, with the similarity of predictions from the *other* neural network as a covariate, to assess relationships between model-based representations of looming and self-reported experience, independent of static visual features, and vice versa.
